# Longitudinal Study on Ocular Manifestations in a Cohort of Patients with Fabry Disease

**DOI:** 10.1101/557207

**Authors:** Langis Michaud

## Abstract

**Purpose:** This study aims to assess the evolution of ocular manifestations in a cohort of Fabry patients.

**METHODS:** This is a prospective observational study conducted from 2013 to 2017 (5 consecutive exams). All subjects underwent a comprehensive ocular examination including oriented case history, refraction, corneal topography, biomechanical corneal properties and pachometry assessments, aberrometry, anterior segment evaluation, double-frequency visual field (FDT), intra-ocular pressure, and ocular fundus. At baseline, 41 subjects enrolled but 9 dropped-out and 4 files were not kept for analysis (missing data). Remaining 28 subjects were classified into: Group 1 -hemizygotes (HMZ), all on enzyme replacement therapy (ERT) (N=10); Group 2 -heterozygotes (HTZ) actively ERT-treated (N=8), and Group 3 -HTZ not treated (N=10).

**RESULTS:** There is a high intra and inter-subjects variability. At baseline, prevalence of the ocular manifestations found is similar to published data: cornea verticillata (89.2%), conjunctival vessels tortuosity (85.7%), corneal haze (67.8%), retinal vessels tortuosity (64.2%), anterior cataract (39.2%) and posterior cataract (28.5%). Prevalence for new elements are found: upper lid vessels toricity (96.4%) and micro-aneurysms (42.8%). At the end, micro-aneurysms (+82%), posterior cataract (+75%) corneal haze (+21%) anterior cataract (+17%) and retinal vessels tortuosities (+4%) evolved in prevalence and severity despite the fact that 68% of the patients were on ERT. Treated heterozygotes evolved more than other groups (p>0.05)

**CONCLUSION:** ERT does not halt the clinical evolution of several ocular manifestations. Longer observational time may be required to fully confirm these findings.

## INTRODUCTION

Fabry is qualified as a pan ethnic X-linked inherited condition and is considered a rare disease(1); however, it represents the second most prevalent of the 50 known lysosomal storage disorders(2), which are all characterized by a cell deposit of a substrate, within lysosomes, as a result of abnormal enzymatic activity. With Fabry, the absence or deficient activity of α-galactosidase A leads to the accumulation of globotriaosylceramide (Gb-3) in a variety of cells (renal, endothelial, cardiac, dorsal root ganglion neural)(3).

As the disorder evolves, and the substrate continues to build-up, cellular dysfunction will trigger organ impairment and eventually system damages, leading to substantial morbidity and reduced life expectancy(4), especially for patients left untreated(5).

The manifestations of the disease are highly variable, depending on genetic mutation and the gender of the patient. Based on mutations, patients can be categorized into 4 groups(6). A first group of GLA mutations is known as null alleles and is associated with an absent or non-functional enzyme protein. This is considered the classic Fabry disease, associated with severe clinical manifestations. A second group is labelled as missense mutations resulting in polypeptides with an amino-acid replacement, probably not leading to a null allele condition but still generating serious clinical negative outcomes. Mild missense and glycosylation mutation p.N215S are defining a third and fourth group of subjects, both characterized with some enzymatic residual activities, and associated with an attenuated phenotype. In the past, asymptomatic subjects were only considered carriers of the disease(7), but must be considered true Fabry patients(8).

Gender divides patients into two distinct groups. Men are defined as hemizygotes, which are individuals having only one allele for a specific characteristic(9). This means that a gene located on the X chromosome has no corresponding gene on the Y. Hemizygotes can only inherit the deficiency (X) from their mother and will automatically transmit the disease to their daughters (boys receiving the normal Y). Females are considered heterozygotes, having different alleles at a particular gene locus on homologous chromosomes(9). They can inherit their defective X from both parents. They have a 50% chance of transmitting the disease to any of their children, regardless of gender.

The clinical portrait becomes more complex when both classifications are combined. This means that hemizygotes and heterozygotes can present little or no residual enzymatic activity, which obviously impacts the severity of the disease and its clinical manifestations. For heterozygote patients, the clinical expression will be highly influenced by lyonization, a phenomenon that occurs when a defective gene expression is countered by a normal one, and consequently expressed to varying degrees. As of 2018, 840 possible mutations were compiled(10), which explains the high variability of clinical findings among patients diagnosed with Fabry.

### Clinical manifestations

In the case of Fabry, the presence of substrate accumulation can be traced back to the prenatal period(11) but generally, symptoms do not develop before early childhood (4-6 years old)(2), boys being affected earlier than girls(12). From then on, affected children may suffer from acute and chronic neuropathic pain, which manifests itself as a burning sensation and tingling in the hands and feet (acroparaesthesiae). They may also show reduced or absent sweating (hypohidrosis), which eventually makes it difficult to exercise(13) and perform physical activities(14). Finally, abdominal pain and postprandial diarrhoea may be reported. Fabry children, mostly boys, exhibit greater weight and height variations, most of them remaining under the normal development curve(15). During adolescence, gastro-intestinal problems may increase in severity and frequency, and proteinuria can become more elevated(2). Delayed puberty and abnormal sexual functions were reported(16). Fabry crisis, manifested as severe pain radiating from the hands and feet, may occur spontaneously as a response to various environmental conditions (heat, cold, stress, fatigue, illness, intense exercise, etc.)(5). Swelling of the affected members (lymphedema) and redness may be present and are predictors of future heart and renal dysfunctions(2).

Vascular skin lesions (angiokeratomas) can appear, clustering around the umbilicus and swim trunk areas(17). Later in life, respiratory problems, cardiac and cerebrovascular manifestations may be manifested. Hearing loss or tinnitus are not uncommon at this strage(18). Late disease will be characterized by life-threatening renal failure (more the case for hemizygotes), stroke and severe cardiac problems (more often for heterozygotes)(19).

### Ocular manifestations

Ocular manifestations are among the first observable signs of the presence of the disease, even before birth(20), and are easily identified through a regular slit lamp examination made by a trained eyecare professional. One prevalent feature is the presence of pigments deposited in the cornea, with a whorl-shaped pattern. This keratopathy is known as cornea verticillata(21). By diffusion from the conjunctival blood vessels, the cornea not being vascularized, pigmented substrates accumulate in a linear fashion at the epithelial basement membrane level and the adjacent anterior stroma, primarily on the lower third of the cornea(22). Over time, it may evolve to reach the upper quadrants. Over 90% of hemizygotes show this type of pigmentation by age of 4, while heterozygotes present this clinical sign later, around the age of 10 on average(23). It was reported that patients from groups 1 and 2 (null allele and missense) are more affected than others, with higher prevalence and severity of the manifestations (6). The level of pigmentation is then associated with the progression and severity of the disease(24).

Less specific, a diffuse corneal haze can also be found in conjunction with pigment deposits(6). Both manifestations are not habitually associated with any loss of visual acuity. Differential diagnosis implies keratopathy related to the intake of medication such as amiodarone, tamoxifen, chlorpromazine, indomethacin and gold salts, just to name a few(25). The main difference is the asymmetric distribution of pigmentations between both eyes in the case of Fabry, vs the more symmetrical mirror image provided when pigments are second to drug deposits. Clear understanding of the medical condition of the patient is obviously essential to establish the differential diagnosis.

Lens opacification is less pathognomonic than corneal verticillata and its prevalence is also lower(26). Fabry’s cataract appears as a sub-capsular opacity along the posterior lens suture lines, very similar to the cortisone-induced cataract or those following head trauma. This type of cataract becomes visually disturbing soon after its onset and patients develop high light sensitivity and reduced visual acuity(27). Hemizygotes can show early signs of cataract by age 20. This type of lens opacification may also be seen in patients with mannossidosis (28) and other lysosomal storage disorders. A second cataract manifestation relates to the presence of a white, wedge-shaped, linear deposit on the anterior sub-capsular area of the lens(21), usually covering all quadrants. They are less associated with visual disturbance and may not be visible under slit lamp examination without pupil mydriasis.

Blood vessel tortuosity appears as a consequence of the alteration of the vessels’ natural architecture by substrate accumulation in the vascular endothelium(29). Hemizygotes are almost all affected by this clinical sign after age 30, while at least 50% of heterozygotes show similar manifestation(30). Tortuosity can be seen in the bulbar conjunctiva, in the retina, and as reported earlier, on the external surface of the upper eyelid, where the presence of aneurysms was found highly related to the presence of the same clinical finding in the bulbar conjunctiva(31). Vessel tortuosity is not associated with functional loss affecting vision; however, Fabry-related vasculopathies (artery or veinule occlusions, choroidal neovascularization) are rarely seen (32, 33), except among patients with more severe systemic involvement. Differential diagnosis should include the presence of fucosidosis(34), or gangliosidosis(35), where similar blood vessel alterations were reported.

Alteration of vascular insufficiency may lead to the development of optic nerve ischemia. Enlargement of the blind spot, and central defects in visual field testing, is related to optic neuropathy probably second to this phenomenon(36). Finally, substrate accumulation in the lachrymal gland fosters eye dryness, experienced by 50% of Fabry patients(37). Recently, bulbar conjunctival lymphangiectasia, a sign of ocular surface irritation, was identified as a new ocular finding, in a cohort of Fabry patients particularly symptomatic of eye dryness(38).

All of these ocular manifestations were reported as single events, on several cohorts of patients, with variable methods of data collection. To our knowledge, no publication evaluated the progression of ocular manifestations over time.

### Treatments

Fortunately, since 2001 in Europe and 2003 in the US, enzyme replacement therapy (ERT) is available to manage the patient’s condition, in addition to other medication needed to stabilize or improve cardiac, renal, gastro-intestinal and other systemic implications(39). In Canada, Fabry patients can access ERT if they meet nationally accepted criteria based mostly on cardiac and renal involvements at the time of diagnosis(40). Once approved, patients are randomly assigned to one of the two drugs available. This targeted use of ERT seems to reduce the risk of adverse outcomes related to Fabry(41). Use of supportive therapies such as aspirin, renin-angiotensin blockade and statins are also considered in ERT-treated subjects(41).

In general, ERT was shown to reduce the GL-3 level in the blood and lessen gastro-intestinal symptoms(42). In some patients treated with ERT, renal function decline is slowed(43) and pain crisis is less severe(44). It is also proven that ERT may significantly improve cardiac disease associated with Fabry, lowering the risk of cardiac death(45); however, it does not seem to lower the risk of stroke(46). It also provides better patient outcomes by preventing serious complications like left ventricular hypertrophy (LVH)(47) or by preserving pulmonary function if the treatment is initiated early(48). This confirms that ERT initiation at a younger age results in a better biochemical response, especially in men with classical Fabry(49).

Two synthetic enzymes (ERT) are currently prescribed: agalsidase α (Replagal^®^; Shire Human Genetic Therapies, Inc. Cambridge, MA; 0.2 mg/kg per infusion) and agalsidase β (Fabrazyme^®^; Genzyme Corporation, Cambridge, MA, 1.0 mg/kg body weight infused every 2 weeks as intravenous infusion). Generally, both drugs are considered equivalent, with no significant difference found in clinical events(50). Both aim to stabilize the patient’s condition, if not improve it, knowing that the damages made on cells, organs and systems are mostly irreversible.

However, a greater biochemical response and an improved ventricular mass was observed with agalsidase β(51). An improved estimated glomerular filtration rate (eGFR) is also associated with the use of agalsidase β after switching drug from agalsidase α formulation (52) A recent Cochrane report confirmed that agalsidase β may be associated to a lower incidence of cerebrovascular events than agalsidase α(53).

Oral medication (chaperone) may be prescribed for a subgroup of patients affected by amenable mutations(54). Migalastat increases alpha-galactosidase-A activity, stabilizes related serum biomarkers, improves cardiac integrity(55), renal function, LVMi, plasma lyso-Gb3, and diarrhea symptoms, especially in susceptible classic Fabry patients(56). Other molecules are under development, like PRX-102 (PEGunigalsidase alfa), a novel PEGylated enzyme expressed in a plant production system(57), and are promising as future alternatives to injections(44).

### Impact on the quality of life

There is a strong relationship between the severity of the symptoms and the patient’s quality of life. During childhood, class attendance may be negatively affected(13). Adults may not be attentive to the nonspecific nature of the symptoms or may not believe the children’s complaints about chronic pain, in the absence of external visible signs. This denial of the condition delays the diagnosis and may have a significant psychological impact, children being perceived as cheaters or liars. Not surprisingly, Fabry patients are eventually more inclined to depression(58), alcoholism and drug dependency. Some will even consider committing suicide during their adolescence(59). Higher severity of the symptoms, especially for men, may limit sports and social activities, often resulting in a sedentary lifestyle, low self-esteem and exclusion(60). This is made even more complicated by chronic fatigue (misdiagnosed as depression), intolerance to physical activities and poor self-perception of health, particularly for women(61). Employment opportunities become limited, making economic stress constant.

There are also multiple side effects associated with the treatment, that is demanding for the patient. Consequently, the positive outcomes from enzyme replacement therapy are weighted by the burden of the treatment. This is why it is not so obvious that the quality of life may be improved with ERT(62). With that in mind, it is important to reiterate the usefulness of slit lamp screening by eyecare practitioners, knowing that ocular manifestations related to Fabry occur early in life. Longevity and quality of life may be improved with a timely diagnosis of the patient’s condition(37).

## OBJECTIVE

This study aims to assess the longitudinal evolution of ocular manifestations in a cohort of Fabry patients.

## METHODS

This is a prospective observational study, conducted in adherence to the tenets of the Declaration of Helsinki. It was approved by the Université de Montreal (UM) review board for experimentation on humans. A majority of participants (80%) were recruited after referral from the only metabolic center managing Fabry patients in the province of Quebec at that time (Hôpital Sacré-Coeur de Montréal). Other patients (20%) were referred from optometric practices in Montreal area. Written consent was obtained for all participants, at enrollment, after they had been fully informed of the goals and procedures of the longitudinal study. The only entrance criteria was to having been diagnosed with Fabry disease. All participants accepted to be seen every year from 2013 (baseline) up to 2017 (5 exams) for a comprehensive eye assessment, subject to their health condition.

A complete description of the clinical procedures was described elsewhere(63). To summarize, a list of clinical testing, made at each annual visit, is described in Table 1. All the testing were done in order. Among these procedures, the following ones were defined as essential components of a standard Fabry ocular assessment : a complete case history screening for signs and symptoms related to the disease, ocular anterior segment evaluation under slit lamp, with emphasis on the cornea and crystalline lens, evaluation of the visual field using double-frequency strategy (FDT), and ocular fundus assessment under pupil dilation. Pupil mydriasis was achieved after instillation of 1 drop of tropicamide 1% and 1 drop of phenylephrine 2.5% in each eye, subject to patient’s contraindications.

**TABLE 1.**
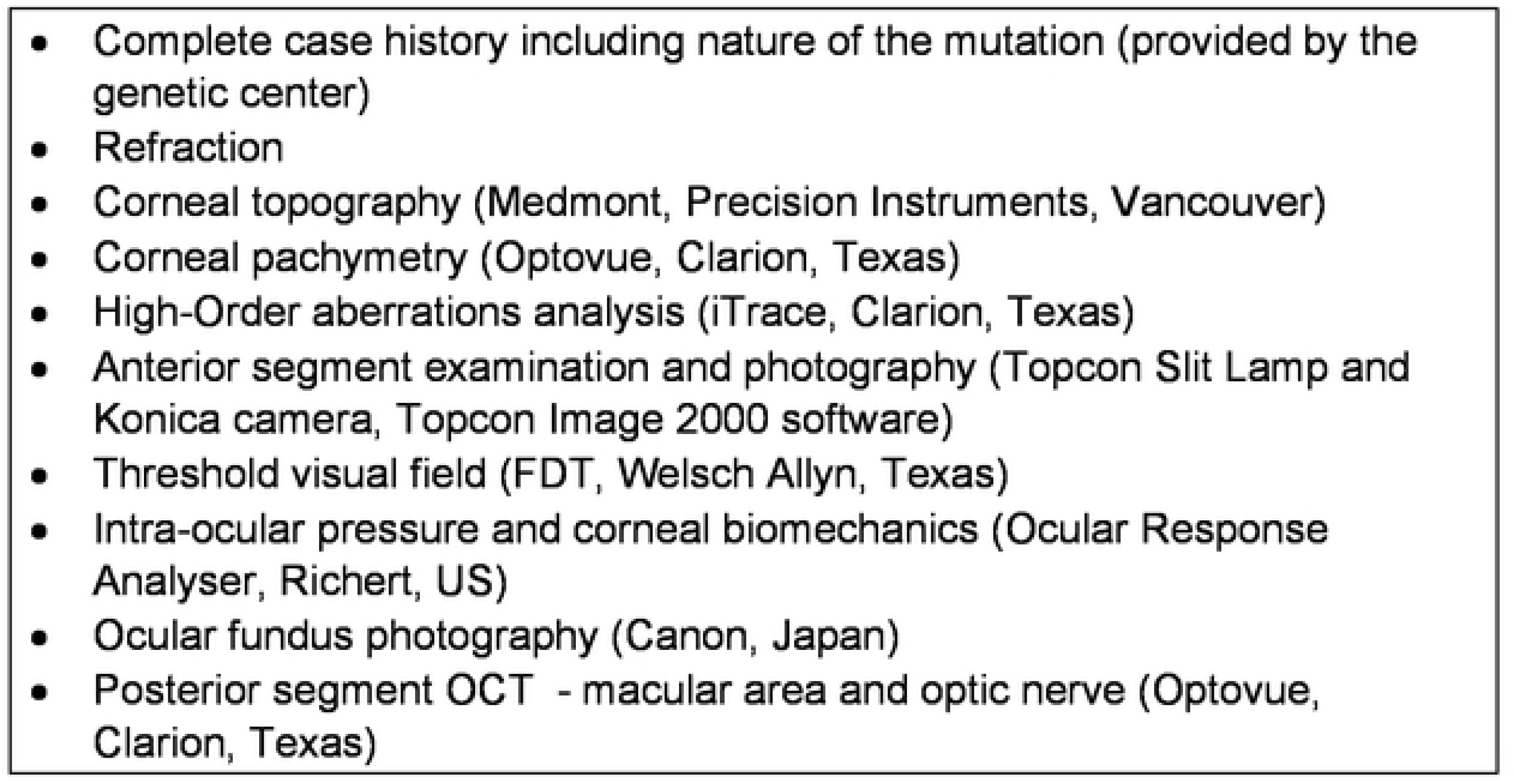
LIST OF TESTING- ANNUAL EXAM

|  |
| --- |
| • Complete case history including nature of the mutation (provided by the genetic center)<br>• Refraction<br>• Corneal topography (Medmont, Precision Instruments, Vancouver)<br>• Corneal pachymetry (Optovue, Clarion, Texas)<br>• High-Order aberrations analysis (iTrace, Clarion, Texas)<br>• Anterior segment examination and photography (Topcon Slit Lamp and Konica camera, Topcon Image 2000 software)<br>• Threshold visual field (FDT, Welsch Allyn, Texas)<br>• Intra-ocular pressure and corneal biomechanics (Ocular Response Analyser, Richert, US)<br>• Ocular fundus photography (Canon, Japan)<br>• Posterior segment OCT - macular area and optic nerve (Optovue, Clarion, Texas) |

The threshold visual field (VF) was tested using frequency doubling technology (FDT). This is a screening strategy based on a flicker illusion that showed high sensitivity and specificity to detect ganglion cells abnormality (leading to glaucoma), but less so for other retinal diseases. In this study, results were classified as normal (no VF defect) or abnormal (VF defect in at least one quadrant). A defect was considered present when a tested point (area) showed a threshold sensitivity (in decibels) reduced by 5% compared to a normal database. (64) Results can be altered by the presence of corneal scars or crystalline lens clinically relevant opacities.

Other testing listed in Table 1 may be considered optional in practice but necessary from a research perspective. One example is the adjunct testing which evaluated the biomechanical properties of the corneas with the use of an Ocular Response Analyzer (ORA; Reichert Ophthalmic Instruments, Depew, NY). The measurements were made by applying a force on the cornea, via a jet of air. Results are based on the ability of the cornea to regain its shape and translates the corneal tissue behaviour. Three parameters were evaluated in this study population: the Corneal Resistance Factor (CRF), the Corneal Hysteresis (CH) and the non-contact corneal-compensated Intra-Ocular Pressure (IOPcc).

Optical aberrations can be defined as distortion in the image formed by an optical system. They can be divided in low-order aberrations, related to the refractive status of the eye, or in high-order aberrations (HOAs), above the previous ones. It is known that these HOAs vary with refractive status, pupil size, presence of corneal or lens opacities, etc(65). HOAs optical impact in the eye differs from one type to another one and is patient dependent(66). Spherical aberration, coma and trefoil are the only ones of clinical interest. Subjects affected with higher refractive errors, and/or those with larger pupils are habitually showing higher levels of aberrations.

HOAs presence in a Fabry’s patient cohort was rarely assessed. In this study, a wavefront aberrometer (iTrace, Tracey Technologies, US) was used. It allows to measure the quality of vision and the visual function using a fundamental thin beam principle of optical ray tracing. This aberrometer sequentially projects 256 near-infrared laser beams into the eye to measure forward aberrations, processing data point-by-point. Whenever a significant finding occurs, it is possible to identify which clinically relevant aberration is most prominent (coma, trefoil, secondary astigmatism or spherical aberrations).

The threshold visual field (VF) was also tested using frequency doubling technology (FDT). This is a screening strategy based on a flicker illusion that showed high sensitivity and specificity to detect ganglion cells abnormality (leading to glaucoma), but less so for other retinal diseases. In this study, results were classified as normal (no VF defect) or abnormal (VF defect in at least one quadrant). A defect was considered present when a tested point (area) showed a threshold sensitivity (in decibels) reduced by 5% compared to a normal database. (64) Results can be altered by the presence of corneal scars or crystalline lens clinically relevant opacities.

All the procedures listed were conducted by the same examiner (author), at the same location (UM clinic) for all participants. The same testing procedure was repeated periodically over five annual consultation and results were compared from baseline to the final visit results.

## STATISTICAL ANALYSIS

Both eyes were considered separately because of the high variability found between them during clinical observations. All calculations were made by an experienced statistician from the UM’s Statistics Department, using version 9.4 of the SAS software, with the 5% significance level.

Each measure was analyzed using a repeated-measure variance analysis with a mixed 4-factor model; these factors include intra-subject factors (time and eye) and inter-subject factors (group and mutation). For the sake of parsimony, given the number of participants and factors studied, the analyses were first run by incorporating all possible terms of interaction. When the triple (3 factors) and quadruple (4 factors) interactions were found to be insignificant, the analyses were redone keeping order 2 interactions. Where no interaction was significant in this last analysis, the latter was redone focusing only on the main effects. For significant interactions, post-hoc analysis were controlled for one factor at a time in order to identify the source of the interactions.

For the categorical variables, the evolution of the initial time at 5 years was categorized according to 3 categories (deterioration, stability and improvement). The association between the evolution categories and the three groups was tested using the exact p-value of the chi-square test.

All results were corrected for multiple comparisons.

## RESULTS

The enrollment began in 2013, defined as baseline, and closed in February 2015. The study ended in March 2017 after the fifth annual exam.

At baseline (2013), 41 subjects composed the study population. In 2017, 9 were lost for follow-up: 2 deceased, 2 moved out of the area and were not reachable, 2 voluntary redrawn, and 3 were not able to travel/be seen because of a poor health condition. Because of partial, unreliable or missing data (< 5 exams completed), 4 other files were not considered for analysis. Consequently, data presented in this report relates to 28 subjects with full dataset.

Subjects are classified into 3 different groups for analysis. Group 1 is composed with hemizygotes (HMZ), all on enzyme replacement therapy ERT (N=10). Group 2 is composed with heterozygotes (HTZ) actively ERT-treated (N=8). Finally, Group 3 is composed with HTZ not treated with enzymes (N=10), because they did not met the criteria set by the CFDI committee. Among those on ERT, agalsidase α and β was given to 8 and 10 subjects respectively at baseline. During the study period, agalsidase β became unavailable for approximatively 2 years and patients were switched over the other formulation. When the medication became again available, four subjects were switched back to their original formulation. At the conclusion of the study, 11/18 (6 HMZ, 5 HTZ) remained on agalsidase α. Analysis of the result is made on the evolution of several parameters, by comparing baseline results with the same ones assessed after 5 annual visits. It was not possible, for administrative reasons, to access to the medical file and to track down the exact time of exposure to each medication for every patient during that time. Statistical analysis will not consider the 2 products used differently.

The most common mutation affecting subjects (11/28) was *p.Ala348Pro*, a rare missense one occurring at position c.1042G>C.(67) Other mutations, and their frequency, are listed in Table 2. For statistical analysis, those single mutations will be considered as a single group (“other mutations”).

**TABLE 2.** LIST OF MUTATIONS AFFECTING AT LEAST 1 SUBJECT OF THE STUDY POPULATION

|  |
| --- |
| • p.Arg220* |
| • p.Arg332aspfs*16 |
| • p.Arg342gln |
| • p.Asn34lysfs*22 |
| • p.Cys12phefs*105 |
| • p.Ex4del? |
| • p.Ex5del? |
| • p.Leu414ser |
| • p.Met187valfs*6 |
| • p.Pro214leu |
| • p.Pro293thr |
| • p.Trp81ser |
| • p.Val254del |

Age of the subjects are reported in Table 3. There is a statistical difference based on the gender, men being younger than females in general (F=6.049; p<0.05), but there is no gender* treatment effect (F=1.440; p>0.05).

**TABLE 3.** AVERAGE AGE OF THE STUDY POPULATION

|  | Average Age (+/- SD)<br>(years old) | Range |
| --- | --- | --- |
| Group 1 (HMZ-ERT) | 37.5 + 9.7 | 23.0 to 52.0 |
| Group 2 (HTZ-ERT) | 56.7 + 9.1 | 43.0 to 70.0 |
| Group 3 (HTZ- not treated) | 52.3 + 15.2 | 35.0 to 77.0 |

### Ocular related parameters

Refraction, pachymetry and ocular biomechanical parameters are reported in Table 4.

**TABLE 4 :** REFRACTION, PACHYMETRY AND CORNEAL BIOMECHANICS

|  | BASELINE |  |  | 5 YEARS |  |  |
| --- | --- | --- | --- | --- | --- | --- |
|  | GROUP 1 | GROUP 2 | GROUP 3 | GROUP 1 | GROUP 2 | GROUP 3 |
|  | HMZ- ERT | HTZ - ERT | HTZ-Non ERT | HMZ- ERT | HTZ - ERT | HTZ-Non ERT |
|  | N= 10 | N=7 | N=11 | N= 10 | N=7 | N=11 |
| Refraction OD (D) | -0.33<br>+0.78 | -1.03 ±<br>2.95 | -2.33 ± 2.96 | Not evaluated |  |  |
| Refraction OS (D) | -0.53 ±<br>0.85 | -0.34 ±<br>3.42 | -2.48 ± 3.05 |  |  |  |
| Pachymetry OD (um) | 546.1 ±<br>35.1 | 522.0 ±<br>52.1 | 541.7 ±<br>27.7 |  |  |  |
| Pachymetry OS (um) | 546.7 ±<br>34.3 | 523.1 ±<br>56.4 | 542.2 ±<br>32.5 |  |  |  |
| IOP OD (mm Hg) | 14.0 ± 2.4 | 14.4 ± 3.6 | 16.0 ± 3.0 | 13.8 ± 3.7 | 16.7 ± 2.1 | 15.7 ± 3.8 |
| IOP OS (mm Hg) | 13.4 ± 2.6 | 14.6 ± 2.6 | 16.2 ± 3.7 | 14.1 ± 3.7 | 15.9 ± 3.9 | 15.5 ± 2.3 |
| CRF OD (mmHg) | 11.1 ± 2.1 | 9.9 ± 1.0 | 10.8 ± 2.2 | 10.2 ± 1.7 | 10.5 ± 1.3 | 10.8 ± 2.3 |
| CRF OS (mmHg) | 11.3 ± 1.6 | 11.2 ± 1.5 | 12.2 ± 3.9 | 10.5 ± 1.6 | 10.5 ± 1.4 | 10.8 ± 2.0 |
| CH OD (mmHg) | 11.1 ± 1.1 | 10.7 ± 0.9 | 10.7 ± 1.8 | 11.1 ± 1.5 | 10.3 ± 1.1 | 11.2 ± 2.0 |
| CH OS (mmHg) | 11.8 ± 1.8 | 11.0 ± 1.6 | 11.6 ± 2.6 | 11.1 ± 1.4 | 10.3 ± 0.9 | 11.2 ± 1.8 |
| HOA OD (um) | 0.167 ±<br>0.036 | 0.230 ±<br>0.094 | 0.195 ±<br>0.128 | 0.135±0.089 | 0.202±0.152 | 0.173 ±<br>0.144 |
| HOA OS (um) | 0.116 ±<br>0.077 | 0.172 ±<br>0.100 | 0.141 ±<br>0.110 | 0.135±0.089 | 0.164±0.101 | 0.157 ±<br>0.223 |

### Refraction

For refraction, the three groups showed low myopia (spherical equivalent) at baseline. The right eye showed no difference based on groups (F=1.779; p>0.05) or mutation (F=1.718; p>0.05). Similar results were observed for the left eye, with no notable differences based on groups (F=1.773; p>0.05) or mutation (F=2.080; p>0.05). This parameter was not assessed every year, refractive error variation being not characteristic of the disease evolution over time.

### Corneal Thickness

Corneal thickness for the right eye was found similar among the 3 groups at baseline (F=1.363; p>0.05). There was no difference based on mutation (F=0.499; p>0.05). Similar findings applied for the left eye in which no difference among groups (F= 1.332; p>0.05) or based on mutation (F=0.385; p>0.05). These results are comparable to the average corneal thickness observed in a Caucasian population (550 um)(68).

### Biomechanical: Corneal hysteresis (CH)

Corneal hysteresis was found highly variable among participants. It varied from a minimum of 10.3 (Group 2, OD and OS at 5 years) to a maximum of 11.8 (Group 1, OS, @ baseline). There was no significant difference among groups at baseline (OD: F=1.79; p>0.05; OS: F=0.4615; p>0.05), and the same finding applied at the end of the study (OD: F=1.41; p>0.05; OS F=7.91; p>0.05). These values were considered comparable to those measured on a non-Fabry population (NFP-10.3 to 11.8 vs 9.8 mm Hg to 10.8 mmHg)(69).

### Biomechanical: Corneal resistance factor (CRF)

In this study, CRF varied from 9.9 + 1.0 mm Hg to 12.2 + 3.9 mm Hg (Group 2 OD and Group 3 OS at baseline). There was a significant CRF difference among the groups at the beginning of the study, considering the right eye (F=4.12; p<0.05 OD; F=0.03; p>0.05 OS), Group 2 showing lower values vs Group 1 and Group 3. Also, this group (2) showed increased values more than Group 1 over time (p<0.05; 95%CI [−2.496, −0.145]), for the right eye again, but those results were not duplicated for the left eye (p>0.05; 95%CI [−1.412, 1.362]). There was also a time*eye*mutation effect (F=4.91; p<0.05), which implies that one eye (OD) of patients with the pAla348Pro mutation showed systematically higher values (stiffer corneas) if compared with those carrying other mutations, and that this difference was confirmed over time. Despite high inter-subject variability, average values reported here about Fabry patient’s CRF were found similar to those of a non-Fabry population (NFP=10.5 mmHg)(70, 71).

### High Order Aberrations (HOA)

At baseline, total HOA varied from 0.116 ± 0.036 um (Group 1, OS) to 0.230 ± 0.094 um (Group 2, OD) with no significant difference among groups (OD F=0.396, p>0.05; OS F=0.138, p>0.05). The same findings are made at the end of the study (OD F=0.326, p>0.05; OS F=0.025, p>0.05). There was no significant evolution (increased or decreased values) over time (OD F=0.426, p>0.05; F=0.336, p>0.05). For reference, values of individual HOAs in the normal eye are randomly scattered around zero and the total RMS average value is 0.330 um(72). This cohort of patients showed lower values, then not generating visual symptoms. Higher HOAs can also be found in the presence of nuclear cataracts(73). In this study, based on the results found, anterior or posterior lens opacities were not developed enough to increase HOAs to a significant level.

### Intra-ocular pressure (IOP)

Intra-ocular pressure adjusted for corneal biomechanical parameters (IOPcc) varied from 13.4 + 2.6 mm Hg (Group 1 OS, at baseline) to 16.7 + 2.1 mm hg (Group 2 OD, after 5 years). There was no significant effect observed among the the 3 groups at baseline and after 5 annual visits (F=2.99; p>0.05 OD; F=2.78; p>0.05 OS). In general, IOPcc remained stable over time (F=1.86; p>0.05 OD; F=0.46; p>0.05OS). IOPcc under 21mm Hg is not considered a risk factor to develop ocular pathology, specifically glaucoma(74).

### Threshold visual fields – FDT

Findings for each group are reported in Table 5. At baseline, 5/10 Group 1 subjects showed abnormal VF, which was also the case for 3/8 Group 2 subjects and 5/10 Group 3 subjects. At the end of the study, there was almost no change in these findings, prevalence remaining the same: 5/10 in Group 1, 4/8 in Group 2 and 7/10 in Group 3 showed abnormal VF findings. Visual fields defects remained stable for OD on 24/28 subjects and 27/28 for OS. For those who varied, the location and intensity of the threshold defect differed from one exam to another. There was no statistical difference in the evolution among the groups (χ^2^=3.7048, p>0.05 OD; χ^2^=1.6027, p>0.05 OS). Obviously, a normal population would not show any abnormal field defect.

**TABLE 5:** VISUAL FIELD FINDINGS

|  | GROUP 1 |  | GROUP 2 |  | GROUP 3 |  | TOTAL |  |
| --- | --- | --- | --- | --- | --- | --- | --- | --- |
|  | OD | OS | OD | OS | OD | OS | OD | OS |
| Baseline<br>(abnormal/normal) | 6/4 | 6/4 | 3 /5 | 4/4 | 5/5 | 6/4 | 14 / 14 | 16 / 12 |
| 5 years<br>(abnormal/normal) | 8/2 | 7/3 | 4/4 | 4/4 | 6/4 | 6/4 | 18 / 10 | 17/ 11 |

### OCULAR MANIFESTATIONS

Anterior segment ocular manifestation findings are summarized in Table 6. All findings were graded according to the Fabry Outcome Survey(75) grading scale(19).

**TABLE 6:**
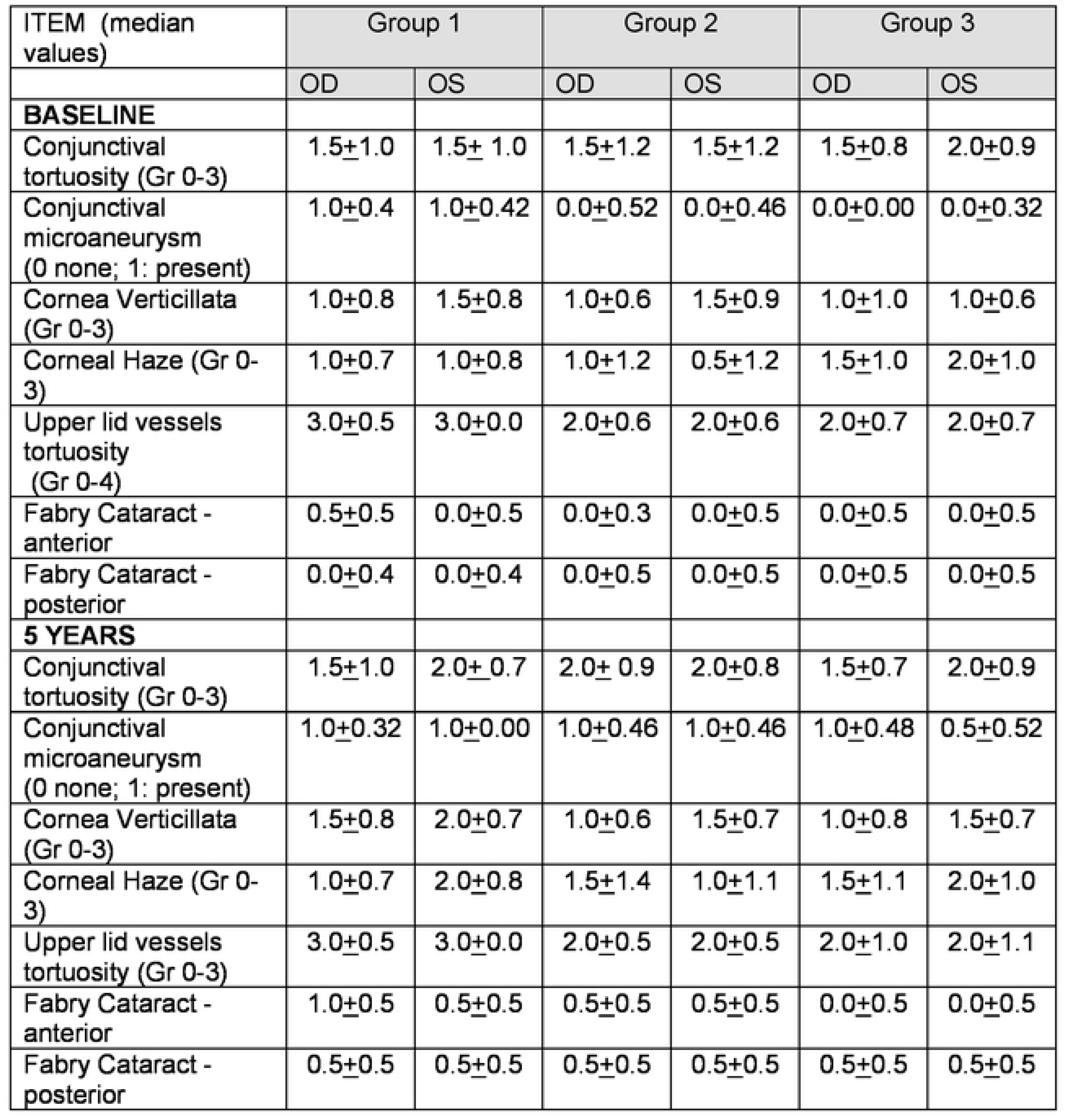
ANTERIOR SEGMENT FINDINGS

### Conjunctival tortuosity

Bulbar conjunctival tortuosity was found in almost every subject at baseline (24/28), except for 1/10 in Group 1, 2/8 in Group 2 and 1/10 in Group 3. At the end, only one subject (Group 2) did not show this finding. On average, tortuosities were evaluated as moderate, varying from grade 1.5 to 2.0 on a scale of 3 (FOS grading scale). Only Group 2 (HTZ-ERT) evolved in severity over time, with a progression from 1.5 to 2.0, but this evolution was not considered statistically significant (p>0.05; 95%CI [-0.136, 0.615] OD p>0.05 95%CI [-0.122, 0.651] OS), most likely because of the low number of subjects. Other variations affecting Group 1 and 3 were not found significant as well (F=0.51; p>0.05 OD; F=0.91; p>0.05 OS); however, there was a time*group*mutation effect (F=5.45; p<0.05) at baseline but not at the end.

### Upper lid vessel tortuosity and microaneurysms

Upper lid vessel tortuosity is a new clinical finding that has been recently identified(31). Again here, almost every subject showed this manifestation, with the exception of one person in Group 3 at the beginning. The same finding applied after 5 annual visits. This makes this ocular manifestation the most prevalent one. Subjects in Group 1 showed more severe vessel tortuosity associated with the presence of microaneurysms (grade 3 out of 4) at baseline (F=9.226; p=<0.05) and at the end (F=3.419; p<0.05), in both eyes, compared to other groups. There was no difference in the evolution in prevalence or severity among groups over time as all remained stable from baseline (F=0.34; p>0.05 OD; F=2.31; p>0.05 OS).

### Conjunctival Microaneurysms

Microaneurysms (MA) were detected mainly in the bulbar conjunctiva, but also in the inner canthus, the lower palpebral conjunctiva, near the lid margin and on the superior external eyelid as well (see 5.2.2.). At baseline, 8/10 subjects of Group 1 showed at least one MA in at least one eye, which was the case for 3/8 subjects in Group 2 and 1/10 in Group 3. This difference was found to be significant (F=3.88; p<0.05 OD; F=3.761; p<0.05 OS), especially considering OD between Groups 1 and 3 (p<0.05; 95%CI [0.14, 0.91] OD); p<0.05; 95%CI [-0.02, 0.85] OS); and at a lesser extent between Groups 2 and 3 (p<0.05; 95%CI [−0.95, −0.14] OD); p<0.05; 95%CI [−0.82, 0.06]OS). At the end, Group 1 remained stable with 9/10 subjects affected, which contrasts with a marked evolution in prevalence for Group 2 (6/8) and Group 3 (7/10). This evolution was found to be significant for the right eye (χ^2^=7.5616, p<0.05) but not for the left eye (χ^2^=1.8667; p>0.05), confirming the high intra-subject variability between both eyes.

Once a MA was found in the bulbar conjunctiva, it was usually found also on the superior external eyelid. This association is present in Group 1 at baseline (5/10) increasing in prevalence with time (7/10), a similar situation occurring at a lesser extent in Group 2 (4/8 at baseline and 5/8 after 5 years). There was almost no association found in Group 3 at baseline (0/10), but this group became more aligned with others at the end of the study, with a higher rate (4/10). This difference among groups was found to be significant (F=3.492; p<0.05 OD; F=4.846; p<0.05 OS); however, the evolution in the numbers of positive association over time was considered the same among the groups (χ^2^= 2.3407; p>0.05 OD; χ^2^= 6.7183; p>0.05 OS).

### Cornea verticillata and haze

Corneal verticillata is certainly considered an important hallmark of ocular manifestations related to Fabry. Corneal pigmentation was found in 10/10 Group 1 subjects at baseline, and at the end. Similar data was observed in Group 3 (9/10 at baseline and after 5 years). In Group 2, subjects slightly evolved in prevalence from 5/8 to 6/8.

As reported in Table 6, Group 1 showed more severe verticillata grading than the other groups at baseline, but this was not considered a significant difference (F= 0.562, p>0.05 OD; F=0.633, p>0.05 OS). This was still the case at the end of the study. All groups in this study evolved in intensity by 0.5 deg (out of 4) over 5 years, which was considered a significant progression if compared to baseline (F=92.153, p<0.05 OD; F=5.817, p<0.05 OS).

Verticillata can be accompanied by corneal haze, which is described as a loss of corneal transparency spreading habituall along the vertical meridian. At baseline, 7/10 subjects in Group 1 showed low level (Grade 1 to 2) of corneal haze, much like 4/8 subjects in Group 2. For Group 3, 8/10 subjects were identified with moderate corneal haze (grade 2 to 3) on average. At the end, all Group 1 subjects displayed corneal haze. Groups 2 and 3 remained stable with 5/8 and 8/10 subjects affected respectively. Overall, groups were found to be statistically similar at baseline (F=0.745; p>0.05, OD; F=1.670; p>0.05 OS). Their evolution varied over time, at least in terms of OS (F=0.08; p>0.05 OD; F=4.14; p< 0.05 OS). More specifically, there was a significant difference in the evolution in prevalence of Group 1 vs Group 2 (p<0.05; 95%CI [0.0624, 1.1207] OS), male patients treated with ERT (HMZ – ERT) showed more severe manifestations with time than female patients on ERT treatments. It is then not surprising to find a time*eye*group*mutation effect (F=3.48; p<0.05).

### Cataracts

Fabry’s cataracts can develop in the anterior portion of the crystalline lens, or can be seen as a sub-cortical posterior lens opacity(37). At baseline, anterior cataract was seen in 5/10 subjects of Group 1, 2/8 of Group 2, and 4/10 of Group 3. Subjects were considered as showing light to moderate opacities (grade 0.5 to 1). This condition remained stable over time except for Group 2, showing a higher prevalence with 4/8 subjects affected at the end of the study. This represents twice the frequency of HTZ-ERT subjects, from baseline, especially marked on the right eye with a significant statistical difference (χ^2^= 5.3375; p<0.05 OD; χ^2^= 2.9120; p>0.05 OS).

Posterior sub-capsular cataract was identifiable, at baseline, on 2/10 subjects of Group 1, 2/8 of Group 2, and 3/10 of Group 3. Grading reveals very early cataract onset. There was no significant difference among the groups (F=0.510; p>0.05 OD; F=0.938; p>0.05 OS). ERT-managed subjects (Group 1 and Group 2) evolved significantly in prevalence over time, ending the study with a doubled number of affected subjects (Group 1: 5/10; Group 2: 4/10). Group 3, females untreated, did not evolve significantly in prevalence, with 4/10 subjects with lens opacification at the end of the study. Severity increased with time, especially for Group 1 individuals where 2 patients were referred for cataract surgery, second to disturbing light sensitivity. None of the other groups underwent surgery; however, most of the patients affected with some degree of sub-capsular posterior opacification reported symptoms of high light sensitivity, making it difficult for them to function in bright light conditions without wearing sunglasses or a hat to generate more shade.

### Retinal vessel tortuosity

Blood vessel tortuosity can be seen in arterioles and veinules. If present, such modifications are graded from 0 (none) to 3 (severe), based on FOS scale(19). Results can be found in Table 7. There were differences among the groups. At baseline, 9/10 subjects showed vessel tortuosity in Group 1, compared to 5/8 in Group 2 and 4/10 in Group 3. The difference was found to be statistically significant (F=3.781; p=0.0481 OD; F=4.634; p=0.0401 OS). More specifically, HTZ-ERT subjects seemed to be more affected (prevalence) than HTZ-Non ERT subjects (p=0.0438; 95%CI [−1.97, −0.41]). There was no significant evolution in prevalence and severity of the condition over study time.

**TABLE 7.**
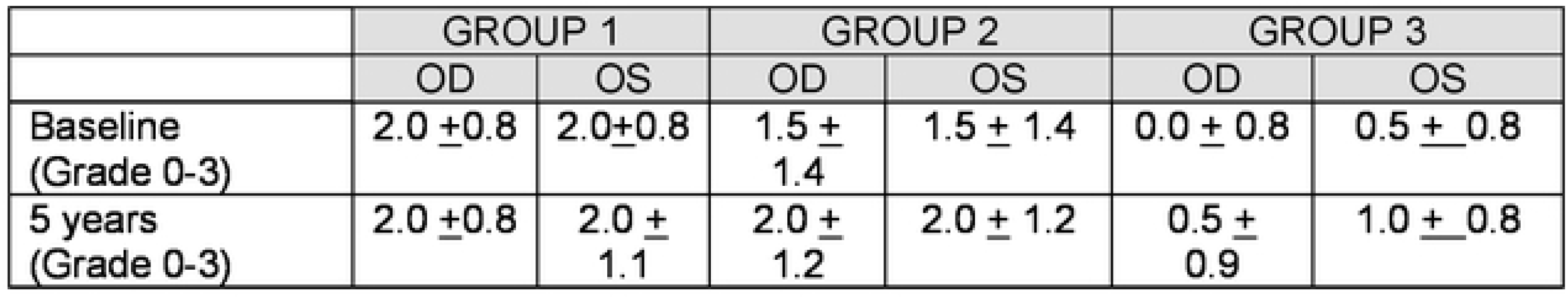
POSTERIOR SEGMENT FINDINGS

|  | GROUP 1 |  | GROUP 2 |  | GROUP 3 |  |
| --- | --- | --- | --- | --- | --- | --- |
|  | OD | OS | OD | OS | OD | OS |
| Baseline<br>(Grade 0-3) | 2.0 ± 0.8 | 2.0 ± 0.8 | 1.5 ± 1.4 | 1.5 ± 1.4 | 0.0 ± 0.8 | 0.5 ± 0.8 |
| 5 years<br>(Grade 0-3) | 2.0 ± 0.8 | 2.0 ± 1.1 | 2.0 ± 1.2 | 2.0 ± 1.2 | 0.5 ± 0.9 | 1.0 ± 0.8 |

It is interesting to note that blood vessels silver wiring was noticed in 2 patients (HMZ- advanced cases in severity), an occurrence that was not reported before.

## DISCUSSION

This study tried to establish the prevalence and the natural course in intensity of ocular manifestations, over 5 annual visits, in Fabry patients. Prevalence of the ocular manifestations found in this study (see Table 8) can be easily compared to the results from a review published earlier(30). All results are in keeping with known prevalence, although variations may occur with different demographic features, subject genotypes, technology used to evaluate clinical manifestations and the subjectivity of the observers. Based on this study results, it is possible to identify the stage of the disease evolution, as another factor to consider. A patient on ERT-treatment is expected to show more signs, with increased severity. Consequently, it is not possible to exclude totally the age effect on the results presented here, all 3 groups being considered older Fabry patients. Another cohort, composed of younger Fabry patients, would show a different clinical picture.

**TABLE 8.**
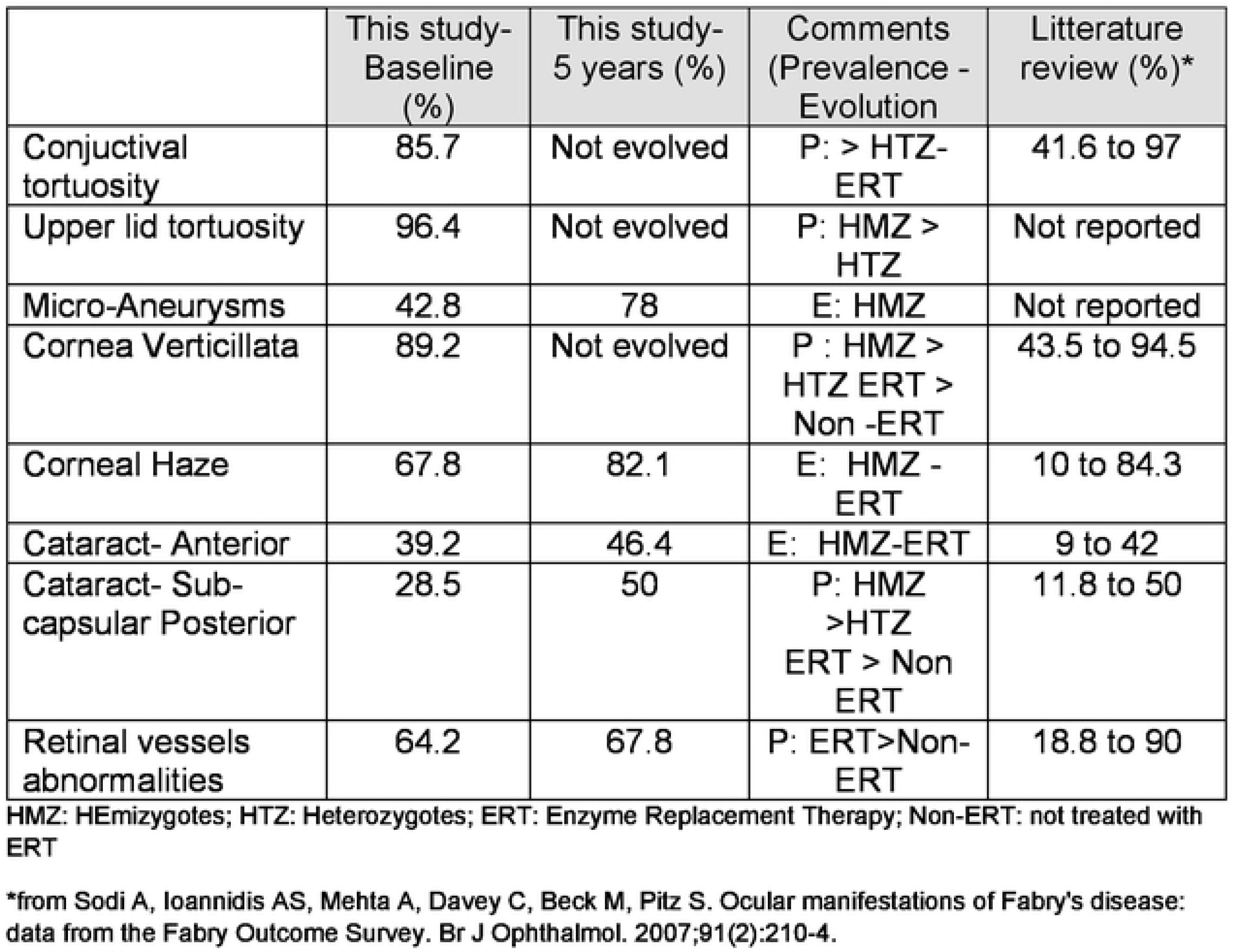
COMPARED PREVALENCE OF OCULAR MANIFESTATIONS

One important finding coming from the analysis of our data (Table 8) is the fact that ERT does not seem to halt the clinical evolution of several ocular manifestations as it does influence positively body’s functions. We did see an increased prevalence in MA (+35%), Corneal Haze (+14%), especially for females treated, anterior cataract (+26%) for males treated; and finally posterior cataracts (+22%), in treated patients overall. Evolution in the prevalence of these signs over the study time was confirmed, despite the regular use of enzyme therapy.

Some would argue that switching drugs during the study may have influenced these results. This was not the case for cardiac function.(76) Looking specifically at the same population studied here, it was reported that switching from alphagal-beta to alfa form, both in standard dose, was associated with a transient increase in clinical events for 6 months but thereafter the clinical event rate returned to baseline for the next 3 years(77). Consequently, it is not likely that the switch of enzyme had influenced the evolution of ocular manifestations observed here.

This study confirmed that cornea verticillata (photo 1) represents a proeminent hallmark feature of Fabry disease, being present in almost every subject (89.2%). Those not showing verticillata were all heterozygotes. Two of them were family related, aged 16 and 48, and affected by p.Arg332Aspfs*16 mutation. Another one (58 y.o.) was showing p.Val254del mutation. None of them were treated (ERT) and their condition was considered less severe, associated with no systemic symptoms. Cornea verticillata, already well developed at baseline in other patients, did not evolve in intensity over time. For not treated females, corneal signs remained stable over time, as their systemic condition.

Corneal haze (photo 2 and 3) is another remarkable feature of Fabry. In this study, some corneas showed haze as the prominent corneal sign over the verticillata. In general, this finding was found in 68% of the patients at baseline, evolving in prevalence to 82% at the study completion. Looking in details, Group 2 (HTZ-ERT) evolve more in prevalence than the others. Putting this in perspective, Group 1 was already advanced, and this confirms that female patients under treatment seemed to tend matching with time the clinical findings of males under treatment.

Pigment accumulation and corneal haze did not affect the level of high-order aberrations nor the biomechanical properties of the cornea. Corneal hysteresis represents the energy absorption during the stress applied on the cornea and is lowered when the cornea is softer or altered by a dystrophy or a structural disease(70). Lower CH values are associated with a higher risk to develop ocular pathology like glaucoma(78). This parameter was never assessed in a Fabry population before. As a reference, CH values measured in a normal cornea population varies from 10.2 to 11.1 mm, depending on the gender, ethnical origin and intra-ocular pressure(79). The study Fabry population showed very similar results. Variations observed may be attributed to ageing instead of being related to the metabolic disorder. It is then possible to conclude that substrate accumulation within the cornea does not modify its biomechanical property.

To the same extent, the corneal resistance factor (CRF) represents the corneal elastic property and is related to the corneal thickness(80). In this study, the CRF varied from 9.9 to 12.2 mm Hg, similar to a non-Fabry Caucasian population, as already mentioned. Again here, variations found are explainable by ageing process instead of being linked with the metabolic disorder itself or any change to the corneal structure second to substrate accumulation.

The presence of micro-aneurysm evolved in prevalence over time. It may suggest a deterioration of the patient systemic condition, more specifically an increased level of substrate accumulation within the blood vessel walls. Usually found in the lower bulbar conjunctiva, MA, in this study, was also found in the inner canthus fornix (see photo 4), at the lid margin (see photo 5), on the external inferior eyelid or superiorly. The variation of the location confirms the fact that ocular manifestation occurs where the substrate accumulates, which is not predictable.

Upper lid vessels tortuosity (see photo 6) was also highly prevalent (96.4%). This is not surprising that the small vessels of the upper lid are showing abnormalities, as they are very similar in structure to those of the bulbar conjunctiva where tortuosities are well documented. Similarity between these 2 areas extend also to the presence of micro-aneurysms on both locations especially among patients ERT-treated. This means that if at least one micro-aneurysm is found either on the bulbar conjunctiva, the inner canthus or the lid margin, it is very likely to find one micro-aneurysm along the vessels or in the area of the external upper lid. The strong association between the occurence of MA on both locations may be used, in the future, as one of the potential validation element for Fabry diagnosis.

It may be surprising that posterior cataract did evolve in prevalence, while anterior cataracts did not change significantly. First of all, posterior cataracts are more prevalent than the anterior one, occurring in up to 70% of the male patients(37), and are known as “classic” Fabry cataracts. Anterior opacities are described as “propeller” and are less frequently seen, depending on where the substrate will accumulate within the body(21). Light sensitivity may be very disturbing and contribute to significantly reduce performance at work or when driving. Other Fabry patients may report visual symptoms including eye dryness, blurry vision, difficulty in seeing under dim or reduced illumination, and soreness/tiredness(75). These symptoms, including light sensitivity, may all be related to lens opacities and should be assessed at every visit. Whenever present, they may prompt cataract surgery on highly symptomatic patients. Evolution of lens opacities may be caused by increased disease severity or simply with ageing. This study was not powered enough to be able to make specific analysis, in that regard, but we may suspect that chances are higher to develop disturbing symptoms as the patient’s condition becomes more severe and the patient is getting older.

Retinal vessel tortuosity (see photo 7) brings another evidence that ERT subjects are more affected than non-ERT subjects. There was a significant difference in prevalence and intensity among the groups, at baseline, which did not vary over time. It is interesting to report that, in 2 individuals (HMZ-ERT) retinal arteries displayed sliver wiring (see photo 8), which was never reported before for Fabry patients. This rare phenomenon is described as a thickening of walls of retinal arterioles leading to the loss of vessels visibility and reflection. This ocular manifestation is habitually related to abnormal systemic blood hypertension but we cannot exclude that, in the case of Fabry, thickening of the walls may be related to high substrate accumulation as a second contributor mechanism. Both subjects were highly symptomatic for Fabry, including a severe cardiac condition, and one died during the course of the study from a cardiac failure.

Visual field defects are manifested as a loss of sensitivity in a point or an area. Approximately 50% of the study population showed such a variation during the study; however, surprisingly, the area where the defect was located at baseline, moved during the following years in a few cases. In order to understand visual field findings, it is important to remember that FDT technology targets magnocellular cells, which are the first ones to be affected by a change in the optic nerve perfusion, which may be altered(81) in the case of Fabry patients, second to GL-3 deposits causing dilatation and instability of the vascular lumen in Fabry patients (21). This result about visual fields cannot be compared to previous findings, where 37% of the patients showed defects manifested as an enlarged blind spot(27). In that case, the Goldman perimetry was used, which implies a seen to unseen strategy for any target presented in the field. Lens opacities (cataracts) may play a role, altering light perception and decreasing sensitivity. However, only 2 patients in this study developed cataracts significant enough to require surgery. Consequently, their influence on the overall results may be considered marginal. Future studies will be needed to confirm these hypotheses.

As a final note, it is important to indicate that, in general, one eye evolved more in prevalence or in intensity of the manifestations, than the fellow eye. This confirms the high asymmetry between the clinical picture found is a vast majority of Fabry patients. If there is one rule about Fabry ocular manifestations, coming from this study, it is that what is found on one eye never mirrors what is seen on the other one.

This study may be limited by the number of drop-outs, expected by the nature of the disease and its evolution over time. However, most of these drop-outs were considered more severe cases. As we saw, the ocular manifestations are more numerous and more intense on such patients. Consequently, our results could have been modified if those subjects would had been able to remain involved and included in the final analysis. To the same extent, the recruitment of a younger population would have modified the findings, assuming that younger patients may be considered less advanced cases, and consequently, showing reduced ocular manifestations in prevalence and intensity.

A second bias is the fact that all males recruited for the study were under treatment. In a perfect world, a group of non-treated hemizygotes would had been followed and compared to treated counterparts, as we were able to do for females. With the same rationale, establishing a correlation between ocular manifestations evolution and other outcomes i.e., kidney, heart, biomarkers that have been reported to respond to ERTs would had been interesting. This study was not designed for this purpose and limited access to the complete file of every patients, for administrative and legal reasons, did not allow to proceed.

A third element to consider is the time of follow-up. For renal outcomes, the CFDI reported a difference after 10 years but not after 5 years(41). Limited funding and drop-out rate over the years reduced the possibility of extending this study.

A fourth bias is the fact that images and clinical data were acquired and analyzed by the same individual. In a perfect scenario, images are taken by one individual and analyzed by a masked observer. Another possibility is to rely on 2 readers to grade clinical findings and to average their score. This was not possible, considering the longitudinal nature of the study and the resources available. Because grading was based on a validated scale (FOS), and because the investigator who analyzed the data was trained to read clinical images, results can be considered as valid.

A fifth element is that, obviously, the results and the conclusion driven from this study can be applied only to a similar population of patients. Extrapolation to other Fabry’s patient cohorts, different in ethnical origin or disease severity, may seems to be logical, but may be hazardous. However, author is confident that this study provides a new understanding on ERT outcome on ocular manifestations and sheds a new lightning on the natural evolution of them over time.

## CONCLUSION

It was possible to establish that five of the known ocular clinical signs related to Fabry did significantly evolve in prevalence and/or in intensity over 5 annual visits, despite enzyme replacement treatment. Micro-aneurysms (+82%) was the clinical manifestation that evolved the most in prevalence, followed by classic (posterior) Fabry’s cataract (+75%) corneal haze (+21%) and propeller (anterior) cataract (+17%). Three other manifestations remain stable over time. At the end, ocular manifestations of ERT-Heterozygotes were comparable with treated hemizygotes (HMZ). Longer observational time may be required to fully confirm these findings.

## Acknoledgements

Dr Daniel Bichet M.D. – UM professor – metabolic disorder clinic Hopital Sacre Cœur.

## FIGURE LEGENDS

Figure (photo) 1: Typical Cornea Verticillata with haze

Figure (photo) 2: Corneal Haze as seen at baseline (1^st^ visit)

Figure (photo) 3: Cornal Haze of the same patient (photo 2) 5 years later, showing increased severity

Figure (photo) 4: Microaneurysm located in the inner canthus (fornix)

Figure (photo) 5: Microaneurysm as seen near the inferior lid margin

Figure (photo) 6: Upper lid tortuosity – grade 4, with micro-aneurysm

Figure (photo) 7: Low retinal vessels tortuosities (arterioles and veinules)

Figure (photo) 8: Silver wiring of retinal arteries

## REFERENCES

1. Fuller M, Meikle PJ, Hopwood JJ. Epidemiology of lysosomal storage diseases: an overview. In: Mehta A, Beck M, Sunder-Plassmann G, editors. Fabry Disease: Perspectives from 5 Years of FOS. Oxford: Oxford PharmaGenesis; 2006.

2. Zarate YA, Hopkin RJ. Fabry’s disease. Lancet. 2008;372(9647):1427–35.

3. Chan B, Adam DN. A Review of Fabry Disease. Skin Therapy Lett. 2018;23(2):4–6.

4. Vedder AC, Linthorst GE, van Breemen MJ, Groener JE, Bemelman FJ, Strijland A, et al. The Dutch Fabry cohort: diversity of clinical manifestations and Gb3 levels. J Inherit Metab Dis. 2007;30(1):68–78.

5. MacDermot KD, Holmes A, Miners AH. Anderson-Fabry disease: clinical manifestations and impact of disease in a cohort of 98 hemizygous males. J Med Genet. 2001;38(11):750–60.

6. Pitz S, Kalkum G, Arash L, Karabul N, Sodi A, Larroque S, et al. Ocular signs correlate well with disease severity and genotype in Fabry disease. PLoS One. 2015;10(3):e0120814.

7. MacDermot KD, Holmes A, Miners AH. Anderson-Fabry disease: clinical manifestations and impact of disease in a cohort of 60 obligate carrier females. J Med Genet. 2001;38(11):769–75.

8. Wang RY, Lelis A, Mirocha J, Wilcox WR. Heterozygous Fabry women are not just carriers, but have a significant burden of disease and impaired quality of life. Genet Med. 2007;9(1):34–45.

9. Miller-Keane. Encyclopedia and Dictionary of Medicine, Nursing, and Allied Health, Seventh Edition. 2003.

10. Ortiz A, Germain DP, Desnick RJ, Politei J, Mauer M, Burlina A, et al. Fabry disease revisited: Management and treatment recommendations for adult patients. Mol Genet Metab. 2018;123(4):416–27.

11. Vedder AC, Strijland A, vd Bergh Weerman MA, Florquin S, Aerts JM, Hollak CE. Manifestations of Fabry disease in placental tissue. J Inherit Metab Dis. 2006;29(1):106–11.

12. El-Abassi R, Singhal D, England JD. Fabry’s disease. J Neurol Sci. 2014;344(1-2):5–19.

13. Ries M, Gupta S, Moore DF, Sachdev V, Quirk JM, Murray GJ, et al. Pediatric Fabry disease. Pediatrics. 2005;115(3):e344–55.

14. Ramaswami U, Parini R, Pintos-Morell G. Natural history and effects of enzyme replacement therapy in children and adolescents with Fabry disease. In: Mehta A, Beck M, Sunder-Plassmann G, editors. Fabry Disease: Perspectives from 5 Years of FOS. Oxford: Oxford PharmaGenesis; 2006.

15. Hopkin RJ, Bissler J, Banikazemi M, Clarke L, Eng CM, Germain DP, et al. Characterization of Fabry disease in 352 pediatric patients in the Fabry Registry. Pediatr Res. 2008;64(5):550–5.

16. Foda MM, Mahmood K, Rasuli P, Dunlap H, Kiruluta G, Schillinger JF. High-flow priapism associated with Fabry’s disease in a child: a case report and review of the literature. Urology. 1996;48(6):949–52.

17. Orteu CH, Jansen T, Lidove O, Jaussaud R, Hughes DA, Pintos-Morell G, et al. Fabry disease and the skin: data from FOS, the Fabry outcome survey. Br J Dermatol. 2007;157(2):331–7.

18. Hegemann S, Hajioff D, Conti G, Beck M, Sunder-Plassmann G, Widmer U, et al. Hearing loss in Fabry disease: data from the Fabry Outcome Survey. Eur J Clin Invest. 2006;36(9):654–62.

19. Mehta A, Ricci R, Widmer U, Dehout F, Garcia de Lorenzo A, Kampmann C, et al. Fabry disease defined: baseline clinical manifestations of 366 patients in the Fabry Outcome Survey. Eur J Clin Invest. 2004;34(3):236–42.

20. Tsutsumi A, Uchida Y, Kanai T, Tsutsumi O, Satoh K, Sakamoto S. Corneal findings in a foetus with Fabry’s disease. Acta Ophthalmol (Copenh). 1984;62(6):923–31.

21. Sher NA, Letson RD, Desnick RJ. The ocular manifestations in Fabry’s disease. Arch Ophthalmol. 1979;97(4):671–6.

22. Mastropasqua L, Nubile M, Lanzini M, Carpineto P, Toto L, Ciancaglini M. Corneal and conjunctival manifestations in Fabry disease: in vivo confocal microscopy study. Am J Ophthalmol. 2006;141(4):709–18.

23. Nguyen TT, Gin T, Nicholls K, Low M, Galanos J, Crawford A. Ophthalmological manifestations of Fabry disease: a survey of patients at the Royal Melbourne Fabry Disease Treatment Centre. Clin Exp Ophthalmol. 2005;33(2):164–8.

24. Kalkum G, Pitz S, Karabul N, Beck M, Pintos-Morell G, Parini R, et al. Paediatric Fabry disease: prognostic significance of ocular changes for disease severity. BMC Ophthalmol. 2016;16(1):202.

25. Bartlett JD. Systemic Drugs affecting the Eye. In: Bartlett JD, editor. Ophthalmic Drugs Facts. 21st ed. St-Louis, MO: Wolters Kluwer Health; 2010. p. 335–42.

26. Sodi A, Ioannidis A, Pitz S. Ophthalmological manifestations of Fabry Disease. In: Mehta A, Beck M, Sunder-Plassman G, editors. Fabry Disease Perspective from 5 years of FOS. Oxford, UK: Oxford PharmaGenesis Ltd.; 2006. p. 249–61.

27. Orssaud C, Dufier J, Germain D. Ocular manifestations in Fabry disease: a survey of 32 hemizygous male patients. Ophthalmic Genet. 2003;24(3):129–39.

28. Arbisser AI, Murphree AL, Garcia CA, Howell RR. Ocular findings in mannosidosis. Am J Ophthalmol. 1976;82(3):465–71.

29. Riegel EM, Pokorny KS, Friedman AH, Suhan J, Ritch RH, Desnick RJ. Ocular pathology of Fabry’s disease in a hemizygous male following renal transplantation. Surv Ophthalmol. 1982;26(5):247–52.

30. Sodi A, Ioannidis AS, Mehta A, Davey C, Beck M, Pitz S. Ocular manifestations of Fabry’s disease: data from the Fabry Outcome Survey. Br J Ophthalmol. 2007;91(2):210–4.

31. Michaud L. Vascular tortuosities of the upper eyelid: a new clinical finding in fabry patient screening. J Ophthalmol. 2013;2013:207573.

32. Sher NA, Reiff W, Letson RD, Desnick RJ. Central retinal artery occlusion complicating Fabry’s disease. Arch Ophthalmol. 1978;96(5):815–7.

33. Sodi A, Bini A, Mignani R, Minuti B, Menchini U. Subfoveal choroidal neovascularization in a patient with Fabry’s disease. Int Ophthalmol. 2009;29(5):435–7.

34. Snodgrass MB. Ocular findings in a case of fucosidosis. Br J Ophthalmol. 1976;60(7):508–11.

35. O’Brien J. Generalized gangliosidosis. J Pediatr. 1969;75(2):167–86.

36. Pitz S, Grube-Einwald K, Renieri G, Reinke J. Subclinical optic neuropathy in Fabry disease. Ophthalmic Genet. 2009;30(4):165–71.

37. Sivley MD. Fabry disease: a review of ophthalmic and systemic manifestations. Optom Vis Sci. 2013;90(2):e63–78.

38. Sivley MD, Wallace EL, Warnock DG, Benjamin WJ. Conjunctival lymphangiectasia associated with classic Fabry disease. Br J Ophthalmol. 2018;102(1):54–8.

39. Schiffmann R, Ries M. Fabry Disease: A Disorder of Childhood Onset. Pediatr Neurol. 2016;64:10–20.

40. Sirrs S, Clarke JT, Bichet DG, Casey R, Lemoine K, Flowerdew G, et al. Baseline characteristics of patients enrolled in the Canadian Fabry Disease Initiative. Mol Genet Metab. 2010;99(4):367–73.

41. Sirrs SM, Bichet DG, Casey R, Clarke JT, Lemoine K, Doucette S, et al. Outcomes of patients treated through the Canadian Fabry disease initiative. Mol Genet Metab. 2014;111(4):499–506.

42. Spada M, Baron R, Elliott PM, Falissard B, Hilz MJ, Monserrat L, et al. The effect of enzyme replacement therapy on clinical outcomes in paediatric patients with Fabry disease - A systematic literature review by a European panel of experts. Mol Genet Metab. 2018.

43. Banikazemi M, Bultas J, Waldek S, Wilcox WR, Whitley CB, McDonald M, et al. Agalsidase-beta therapy for advanced Fabry disease: a randomized trial. Ann Intern Med. 2007;146(2):77–86.

44. Schiffmann R, Floeter MK, Dambrosia JM, Gupta S, Moore DF, Sharabi Y, et al. Enzyme replacement therapy improves peripheral nerve and sweat function in Fabry disease. Muscle Nerve. 2003;28(6):703–10.

45. Rombach SM, Smid BE, Linthorst GE, Dijkgraaf MG, Hollak CE. Natural course of Fabry disease and the effectiveness of enzyme replacement therapy: a systematic review and meta-analysis: effectiveness of ERT in different disease stages. J Inherit Metab Dis. 2014;37(3):341–52.

46. Buechner S, Moretti M, Burlina AP, Cei G, Manara R, Ricci R, et al. Central nervous system involvement in Anderson-Fabry disease: a clinical and MRI retrospective study. J Neurol Neurosurg Psychiatry. 2008;79(11):1249–54.

47. Hongo K, Ito K, Date T, Anan I, Inoue Y, Morimoto S, et al. The beneficial effects of long-term enzyme replacement therapy on cardiac involvement in Japanese Fabry patients. Mol Genet Metab. 2018;124(2):143–51.

48. Franzen D, Haile SR, Kasper DC, Mechtler TP, Flammer AJ, Krayenbuhl PA, et al. Pulmonary involvement in Fabry disease: effect of plasma globotriaosylsphingosine and time to initiation of enzyme replacement therapy. BMJ Open Respir Res. 2018;5(1):e000277.

49. Arends M, Wijburg FA, Wanner C, Vaz FM, van Kuilenburg ABP, Hughes DA, et al. Favourable effect of early versus late start of enzyme replacement therapy on plasma globotriaosylsphingosine levels in men with classical Fabry disease. Mol Genet Metab. 2017;121(2):157–61.

50. SSIEM 2016 Annual Symposium - Abstracts : Rome, Italy, September 2016. J Inherit Metab Dis. 2016;39 Suppl 1:35–284.

51. Arends M, Biegstraaten M, Wanner C, Sirrs S, Mehta A, Elliott PM, et al. Agalsidase alfa versus agalsidase beta for the treatment of Fabry disease: an international cohort study. J Med Genet. 2018;55(5):351–8.

52. Kramer J, Lenders M, Canaan-Kuhl S, Nordbeck P, Uceyler N, Blaschke D, et al. Fabry disease under enzyme replacement therapy-new insights in efficacy of different dosages. Nephrology, dialysis, transplantation : official publication of the European Dialysis and Transplant Association - European Renal Association. 2018;33(8):1362–72.

53. El Dib R, Gomaa H, Ortiz A, Politei J, Kapoor A, Barreto F. Enzyme replacement therapy for Anderson-Fabry disease: A complementary overview of a Cochrane publication through a linear regression and a pooled analysis of proportions from cohort studies. PLoS One. 2017;12(3):e0173358.

54. Lopez Rodriguez M. Treatment in Fabry disease. Rev Clin Esp. 2018.

55. Muntze J, Gensler D, Maniuc O, Liu D, Cairns T, Oder D, et al. Oral chaperone therapy migalastat for treating Fabry disease: enzymatic response and serum biomarker changes after one year. Clinical pharmacology and therapeutics. 2018.

56. Germain D, Giugliani R, Bichet DG, Wilcox WR, al e. Efficacy of migalastat in a cohort of male patients with the classical form of Fabry disease in a phase 3 study. J Inherit Metab Dis. 2016;39:S219.

57. Schiffmann R, Huges D, Giraldo P, Gonzalez D, al. e. Novel treatment for Fabry disease—IV administration of plant derived alpha-gal-a enzyme safety and efficacy, 1 year experience. J Inherit Metab Dis. 2016;39:S51.

58. Cole AL, Lee PJ, Hughes DA, Deegan PB, Waldek S, Lachmann RH. Depression in adults with Fabry disease: a common and under-diagnosed problem. J Inherit Metab Dis. 2007;30(6):943–51.

59. Desnick RJ, Brady RO. Fabry disease in childhood. J Pediatr. 2004;144(5 Suppl):S20–6.

60. Gold KF, Pastores GM, Botteman MF, Yeh JM, Sweeney S, Aliski W, et al. Quality of life of patients with Fabry disease. Qual Life Res. 2002;11(4):317–27.

61. Street NJ, Yi MS, Bailey LA, Hopkin RJ. Comparison of health-related quality of life between heterozygous women with Fabry disease, a healthy control population, and patients with other chronic disease. Genet Med. 2006;8(6):346–53.

62. Arends M, Korver S, Hughes DA, Mehta A, Hollak CEM, Biegstraaten M. Phenotype, disease severity and pain are major determinants of quality of life in Fabry disease: results from a large multicenter cohort study. J Inherit Metab Dis. 2018;41(1):141–9.

63. Michaud L, Aurey-Brais C. Improved ways to screen for patients with Fabry disease, involving optometry in a multidisciplinary approach. Can J Optom 2012;74(4):25–32.

64. Landers J, Sharma A, Goldberg I, Graham S. A comparison of perimetric results with the Medmont and Humphrey perimeters. Br J Ophthalmol. 2003;87(6):690–4.

65. Hartwig A, Atchison DA. Analysis of higher-order aberrations in a large clinical population. Invest Ophthalmol Vis Sci. 2012;53(12):7862–70.

66. Lombardo M, Lombardo G. Wave aberration of human eyes and new descriptors of image optical quality and visual performance. J Cataract Refract Surg. 2010;36(2):313–31.

67. Auer-Grumbach M, Toegel S, Schabhuttl M, Weinmann D, Chiari C, Bennett DLH, et al. Rare Variants in MME, Encoding Metalloprotease Neprilysin, Are Linked to Late-Onset Autosomal-Dominant Axonal Polyneuropathies. American journal of human genetics. 2016;99(3):607–23.

68. Nemesure B, Wu SY, Hennis A, Leske MC. Corneal thickness and intraocular pressure in the Barbados eye studies. Arch Ophthalmol. 2003;121(2):240–4.

69. Ayala M. Corneal hysteresis in normal subjects and in patients with primary open-angle glaucoma and pseudoexfoliation glaucoma. Ophthalmic research. 2011;46(4):187–91.

70. Ortiz D, Pinero D, Shabayek MH, Arnalich-Montiel F, Alio JL. Corneal biomechanical properties in normal, post-laser in situ keratomileusis, and keratoconic eyes. J Cataract Refract Surg. 2007;33(8):1371–5.

71. Plakitsi A, O’Donnell C, Miranda MA, Charman WN, Radhakrishnan H. Corneal biomechanical properties measured with the Ocular Response Analyser in a myopic population. Ophthalmic Physiol Opt. 2011;31(4):404–12.

72. Salmon TO, van de Pol C. Normal-eye Zernike coefficients and root-mean-square wavefront errors. J Cataract Refract Surg. 2006;32(12):2064–74.

73. Hashemi H, Khabazkhoob M, Jafarzadehpur E, Yekta A, Emamian MH, Shariati M, et al. Higher order aberrations in a normal adult population. J Curr Ophthalmol. 2015;27(3-4):115–24.

74. Canadian Ophthalmological Society evidence-based clinical practice guidelines for the management of glaucoma in the adult eye. Can J Ophthalmol. 2009;44 Suppl 1:S7–93.

75. Romero-Rangel T, Stavrou P, Cotter J, Rosenthal P, Baltatzis S, Foster CS. Gas-permeable scleral contact lens therapy in ocular surface disease. Am J Ophthalmol 2000;130(1):25–32.

76. Ripeau D, Amartino H, Cedrolla M, Urtiaga L, Urdaneta B, Cano M, et al. Switch from agalsidase beta to agalsidase alfa in the enzyme replacement therapy of patients with Fabry disease in Latin America. Medicina. 2017;77(3):173–9.

77. Morel C, Bichet DG, Casey R, al e. Switch of enzyme replacement therapy (ERT) in the Canadian Fabry Disease Initiative Study (CFDI): Intermediate follow-up at 3 and a half years. J Inherit Metab Dis. 2016;39:S214.

78. Abitbol O, Bouden J, Doan S, Hoang-Xuan T, Gatinel D. Corneal hysteresis measured with the Ocular Response Analyzer in normal and glaucomatous eyes. Acta Ophthalmol. 2010;88(1):116–9.

79. Al-Arfaj K, Yassin SA, Al-Dairi W, Al-Shamlan F, Al-Jindan M. Corneal biomechanics in normal Saudi individuals. Saudi J Ophthalmol. 2016;30(3):180–4.

80. Fontes BM, Ambrosio R, Jr., Jardim D, Velarde GC, Nose W. Corneal biomechanical metrics and anterior segment parameters in mild keratoconus. Ophthalmology. 2010;117(4):673–9.

81. Diaconu V, Michaud L, Forcier P, Garon M. Optic Nerve capillaries blood oxygen investigation in Fabry’s disease patients. Inv Opht Vis Sci. 2015;56(7):2674.

